# hypeR-GEM: connecting metabolite signatures to enzyme-coding genes via genome-scale metabolic models

**DOI:** 10.64898/2025.12.08.692998

**Authors:** Ziwei Huang, Paola Sebastiani, Daniel Segrè, Stefano Monti

## Abstract

Enrichment analysis is a cornerstone of “omics” data interpretation, enabling researchers to connect analysis results to biological processes and generate testable hypotheses. While well-established tools exist for transcriptomics and other omics layers, the development of robust enrichment resources for metabolomics remains comparatively limited. To address this gap, we developed *hypeR-GEM*, a methodology and associated R package that adapts gene set enrichment analysis to metabolomics. *hypeR-GEM* leverages genome-scale metabolic models (GEMs) to infer reaction-based links between metabolites and enzyme-coding genes, enabling the mapping of metabolite signatures to gene signatures and their subsequent annotation via gene set enrichment analysis. We validated *hypeR-GEM* using paired metabolomics-proteomics and metabolomics-transcriptomics datasets by assessing whether genes mapped from metabolites significantly overlapped with differentially expressed proteins or transcripts. We further evaluated whether pathways enriched via *hypeR-GEM*-mapped genes corresponded to those derived from paired proteomic or transcriptomic data. In most datasets analyzed, both the predicted enzyme-coding genes and the associated enriched pathways showed significant concordance with independently derived omics signatures, supporting the utility and robustness of *hypeR-GEM*. Finally, we applied *hypeR-GEM* to the analysis of age-associated metabolic signatures from the New England Centenarian Study. The results revealed consistent enrichment of lipid-related pathways, aligning with the well-established role of lipid metabolism in aging, and highlighted additional pathways not captured in the metabolites’ annotation, demonstrating *hypeR-GEM*’s practical utility in a real-world use case.

## INTRODUCTION

Metabolomics is the comprehensive study of metabolites in cells, tissues, or organisms, integrating advanced analytical technologies with statistical and computational tools to identify and quantify these molecules and extract biologically meaningful information ^1,2^. A critical step in interpreting high-dimensional omics data is *enrichment analysis*, which reduces data complexity and enhances interpretability by identifying biologically relevant processes ^3,4^. While tools for enrichment analysis have rapidly expanded for transcriptomics and proteomics, the availability of suitable tools for metabolomics remains relatively scarce^4–6^. Several metabolomic-specific platforms, such as MPEA^7^ and MBRole^6^, provide metabolite-based enrichment analysis but lack capabilities for multi-omics integration. Statistical frameworks like Metabox^8^, Metascape^9^, and 3Omics^10^ address the integration of transcriptomic, proteomics, metabolomic data within metabolic pathway contexts and offer joint visualization functionalities; however, the enrichment analyses remain metabolite-centric and are restricted to a limited set of predefined metabolite pathways. MetaExplore^11^ facilitates the multi-omics integration by leveraging genome-scale metabolic models (GEMs) to design an automated workflow for the customization and curation of GEMs based on user-defined metabolite inputs. However, it does not support metabolite-to-gene mapping and gene-based enrichment analysis. MetaboAnalyst^12,13^ supports enrichment analyses across multiple omics layers, but only when users also provide transcriptomic or proteomic data, and each layer is analyzed independently. Although it provides joint visualization when multi-omics are available, it cannot infer gene- or protein-level information when only metabolomics data are provided. Overall, most of the existing metabolomics-focused frameworks perform enrichment analyses directly in the metabolite space. While this approach has useful applications, it does not support the connection to the extensive gene-centered biological knowledge and functional annotations available for transcriptomics and proteomics, which would enable a more comprehensive multi-omics integration.

Gene set enrichment based on over-representation analysis (ORA) is a widely used framework for interpreting transcriptomics data, supported by a large suite of analytical tools and curated molecular signature databases derived from existing biological knowledge^14–16^. This framework has also been successfully extended to the analysis of proteomics data^17^. However, applying gene-based ORA to metabolomics remains challenging due to the lack of well-defined and consistent associations between metabolites and genes^5,18,19^.

To bridge this gap, we developed *hypeR-GEM*, an R package and methodology that harnesses the rich connectivity encoded in genome-scale metabolic models (GEMs). GEMs are curated resources that capture all known metabolic components and interactions within a biological system, including metabolites, genes, enzymes, and reactions^20–23^. By leveraging the gene–reaction–metabolite associations within GEMs, hypeR-GEM establishes reaction-based mapping between metabolites and enzyme-coding genes, enabling ORA to be applied to metabolomics data through predicted gene-level annotations. Notably, GEMs encode comprehensive metabolic information that captures the complex interdependencies among genes, proteins, and metabolites, providing a multidirectional platform for connecting different omics layers. For example, Richelle et al.^24^ demonstrated the utility of GEMs in the opposite direction—inferring metabolic task activity directly from transcriptomic data. Similarly, Zheng et al.^25^ used GEMs to uncover biologically meaningful metabolite-mediated cell-cell communication from single-cell RNA-seq data, while Wagner et al.^26^ leveraged GEMs to characterize cellular metabolic states at single-cell resolution. These studies highlight the versatility of GEMs for integrating and interpreting multi-omics data across diverse biological contexts.

We evaluated the performance of hypeR-GEM using paired metabolomics-proteomics and metabolomics-transcriptomics datasets. Specifically, we tested whether enzyme-coding genes mapped from differential metabolomics signatures significantly overlapped with differentially expressed proteins or genes in the corresponding proteomics/transcriptomics datasets. We further assessed whether pathways enriched via ORA using hypeR-GEM-mapped genes aligned with pathways enriched in the signatures derived from the proteomic or transcriptomic data.

Our results demonstrate that, across multiple datasets, both the predicted genes and their enriched pathways show significant overlap with direct proteomic/transcriptomic signatures and their pathway enrichments. These findings validate hypeR-GEM as an effective tool for extending gene-centric enrichment analysis to datasets where only the metabolomics layer is available.

We conclude with a case study where we show the results of applying hypeR-GEM to age-associated metabolomics signatures from the New England Centenarian Study (NECS) (manuscript in preparation). This analysis revealed connections between enriched lipid-related pathways identified in metabolite-space – such as lipid and fatty acid metabolism – and those identified in gene-space via hypeR-GEM-mapped genes, including retinol, tryptophan, and bile acid metabolism. These findings illustrate hypeR-GEM’s ability to uncover previously validated biological mechanisms associated with aging.

## RESULTS

### Algorithm Overview

The hypeR-GEM algorithm employs a reaction-centric strategy to establish systematic connections between metabolites and enzyme-coding genes by leveraging the metabolic relationships encoded in GEMs (see Figure 1). Within a GEM, each metabolite is associated with one or more reactions, in which it participates as either a reactant or a product. Each reaction, in turn, is catalyzed by one or more enzyme-coding genes, with exceptions such as exchange reactions and certain transport reactions that are not catalyzed by enzymes. This metabolite–reaction–gene triad provides a natural framework for mapping metabolites to genes. Specifically, a metabolite is associated to an enzyme-coding gene if the gene encodes an enzyme that catalyzes one or more reactions in which the metabolite is involved. In this mapping step, all metabolites, including currency metabolites such as ATP, NADH, H O, and H, are treated equivalently.

**Figure 1:**
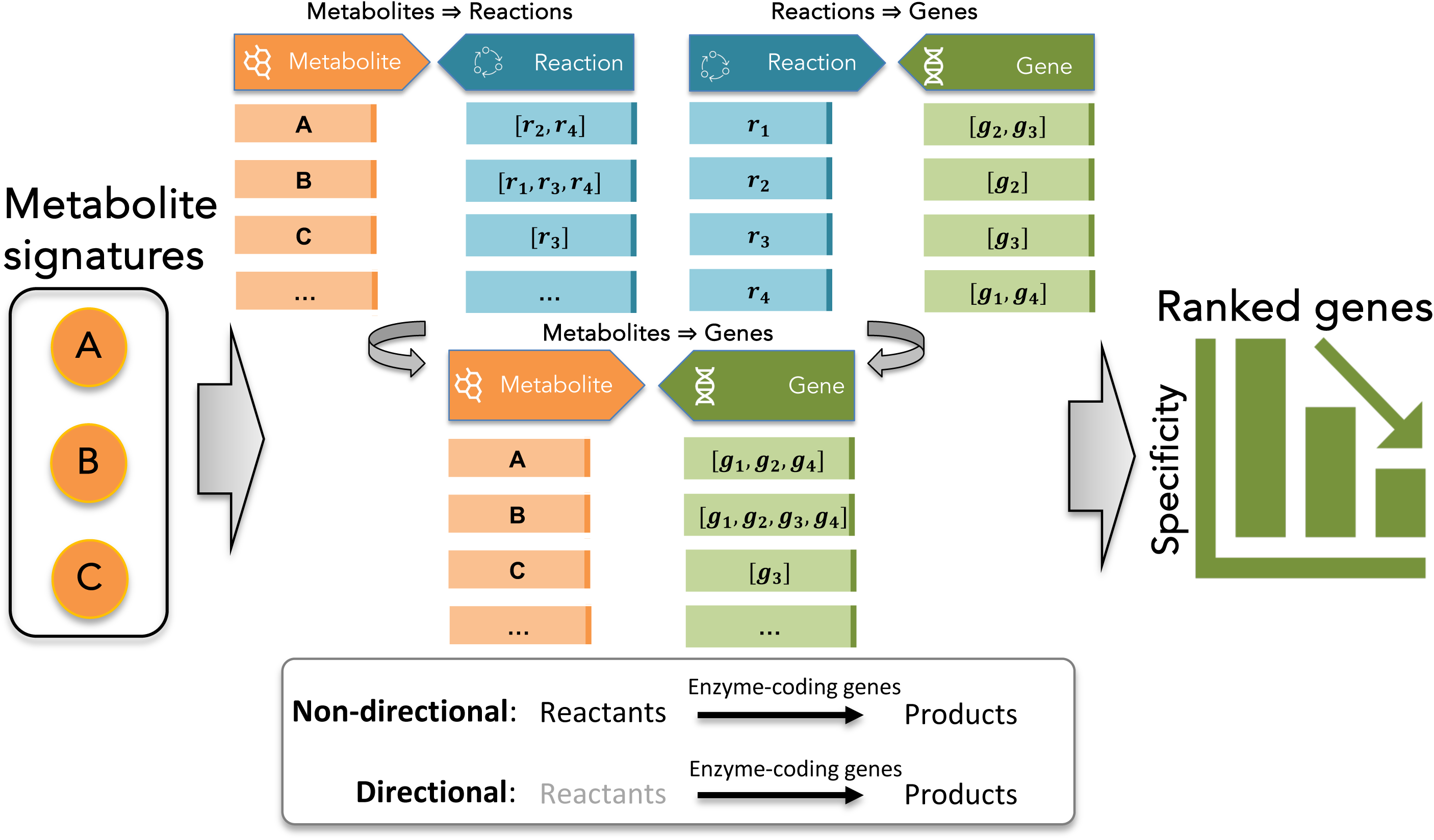
Overview of the hypeR-GEM algorithm. In the schematic, metabolites are denoted by capital letters (e.g., *A*, *B*), metabolic reactions by lowercase *r* with subscripts (e.g., *r_1_, r_2_*), and enzyme-coding genes by lowercase with subscripts (e.g., *g_1_*, *g_2_*). Given an input metabolite signature, hypeR-GEM identifies all reactions in which each metabolite participates and the corresponding enzyme-coding genes that catalyze these reactions. Two mapping strategies are supported: (1) non-directional mapping, linking metabolites through all associated reactions, and (2) directional mapping, restricting associations to reactions in which the metabolite acts as a product. Each mapped gene is subsequently evaluated based on a gene-specificity hypergeometric test to assess the significance of its association with the input signature. The algorithm outputs a ranked list of enzyme-coding genes ordered by their association specificity.

To accommodate different biological contexts, hypeR-GEM supports both non-directional and directional mapping rules, which respectively connect metabolites through all reactions in which they participate or only through reactions in which they are produced, according to the stoichiometry encoded in the model^27^(see “Materials and Methods” for full details). Each mapped gene is subsequently evaluated using a gene-specificity hypergeometric test to quantify the significance of its association with the input signature (see “Materials and Methods” for full details). The final output is a ranked list of enzyme-coding genes ordered by their association specificity as shown in Figure 1.

### Evaluation

We evaluated the performance of hypeR-GEM using paired metabolomics-proteomics and metabolomics-transcriptomics datasets from ten independent studies: (1) TGFβ-induced epithelial-to-mesenchymal transition (EMT) in MCF10A cells ^28^; (2) Serine starvation in HSC3 cells^29^; (3) The New England Centenarian Study (NECS)^30^; (4) - (7) Mouse kidney, liver, gastrocnemius muscle, and plasma from the M005 dataset ^31^; (8) The Religious Orders Study and Memory and Aging Project Study (ROSMAP)^32,33^, and (9) – (10) Urine and serum samples from healthy and COVID-19 patients ^34^ (see “Materials and Methods” for full details).

For each study, we conducted independent differential analyses on one or more phenotypes of interests using linear regression on both metabolomics and proteomics/transcriptomics data to identify molecular signatures (i.e., differentially expressed metabolites, proteins, or transcripts). In total, we analyzed 11 phenotypes across 10 datasets, covering four specimen categories (see Table 4). The metabolite signatures served as input for hypeR-GEM, which mapped them to enzyme-coding genes. Differential analysis and mapping results are provided in Supplementary Data S1-S11. The schematic flowchart of the method performance evaluation is shown in Figure 2. We assessed performance at two levels:

**Figure 2.**
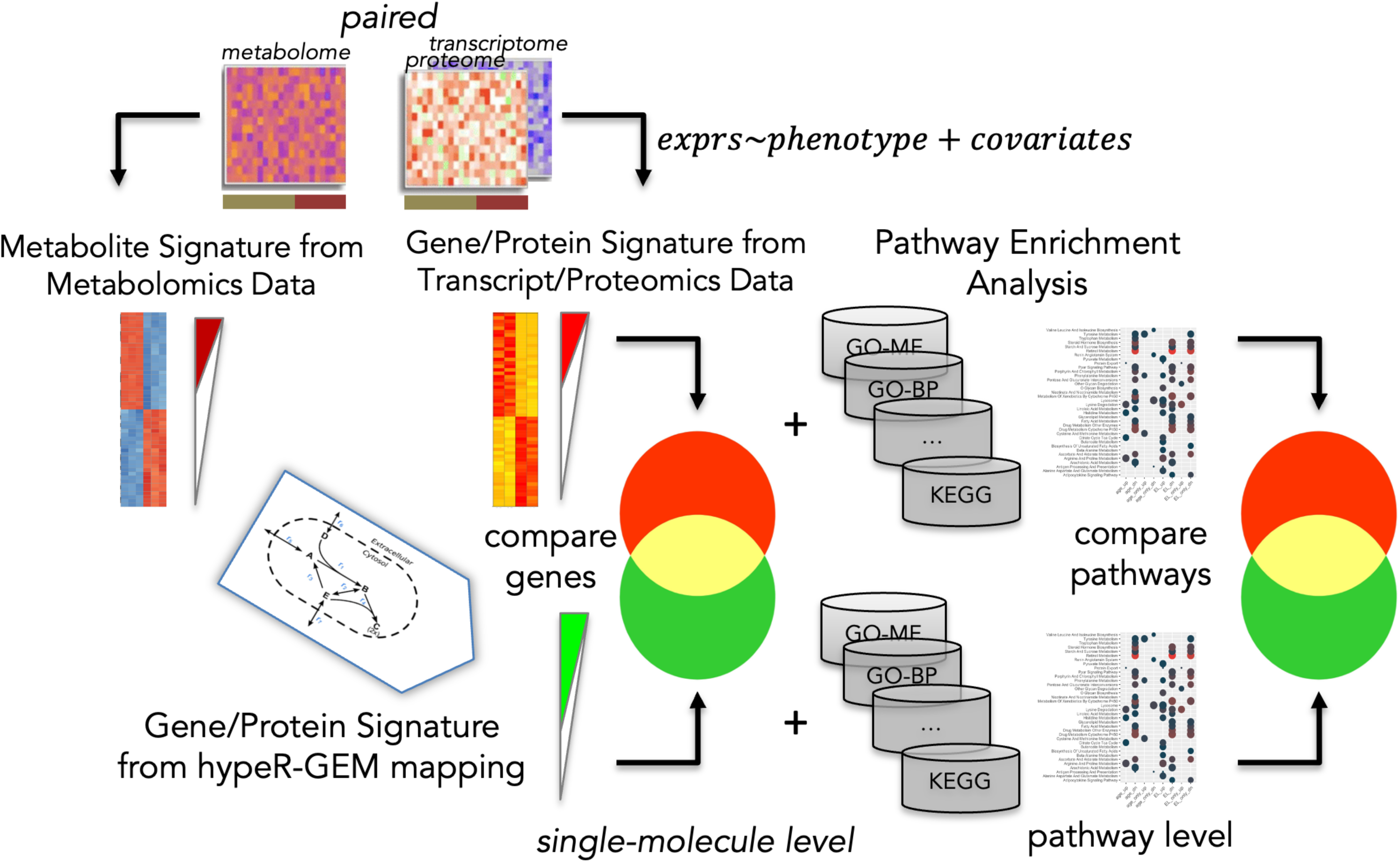
Schematic of method performance evaluation at single-molecule and pathway level. Differential analysis is independently performed on the paired metabolomics and transcriptomics or proteomics datasets to identify molecular signatures. Metabolite signatures are mapped to enzyme-coding genes via hypeR-GEM. At the single-molecule level, we evaluated the significance of the overlap between hypeR-GEM-mapped enzyme-coding genes (green set) and differentially expressed genes or proteins from the paired omics layer (red set). At the pathway level, hypeR-GEM-mapped genes and the corresponding transcriptomic/proteomic signatures are independently annotated by ORA analysis, and the significance of the overlap between their enriched pathways is assessed.

#### Single-molecule level

We evaluated whether the mapped/predicted enzyme-coding genes significantly overlapped with the differentially expressed genes or proteins from the paired omics layer (see “Materials and Methods” for full details). Only comparisons where both omics layers yielded ≥ 10 significant molecules (with mapping in Human-GEM) were considered, as shown in Table 1.

**Table 1.** Evaluation of hypeR-GEM’s performance at single-molecule level. *DE Metabolites*: Number of differentially expressed metabolites in the metabolomics data. *hypeR-GEM (ND/D):* Number of hypeR-GEM mapped enzyme-coding genes based on non-directional (ND) and directional (D) mapping. *DE Genes*: Number of differentially expressed genes/proteins in the transcriptomics/proteomics data. *Overlap (ND/D)*: The overlap between “hypeR-GEM” and “DE Genes”. *Pvalue*: The p-value of the hypergeometric test. *Background*: The background parameter of the hypergeometric test, defined as the intersection between the proteins represented in GEM and those in the corresponding proteomic or transcriptomic dataset.

| **Dataset** | **DE Metabolites** | **hypeR-GEM**<br>**(ND/D)** | **DE Genes** | **Overlap**<br>**(ND/D)** | **Pvalue**<br>**(ND/D)** | **Background** |
| --- | --- | --- | --- | --- | --- | --- |
| *EMT* | 169 | 211/175 | 698 | 124/107 | 6.77e-3/1.99e-3 | 1376 |
| *Serine Starvation* | 36 | 672/577 | 833 | 273/233 | 2.14e-3/7.63e-3 | 2309 |
| *NECS Age* | 291 | 118/98 | 142 | 33/28 | 1.10e-2/1.45e-5 | 722 |
| *NECS EL* | 274 | 132/102 | 142 | 34/26 | 3.63e-2/7.45e-2 | 722 |
| *M005 Kidney* | 328 | 331 /271 | 150 | 50/42 | 7.17e-4/1.34e-4 | 1482 |
| *M005 Liver* | 227 | 322/275 | 402 | 122/108 | 4.50e-2/2.27e-2 | 1291 |
| *M005 Gatroc* | 227 | 74/49 | 28 | 2/2 | 7.76e-1/5.49e-1 | 762 |
| *M005 Plasma* | 143 | 40/34 | 55 | 20/15 | 1.04e-2/1.04e-1 | 164 |
| *ROSMAP* | 61 | 169/128 | 152 | 23/11 | 2.99e-2/6.46e-1 | 1652 |
| *COVID Urine* | 367 | 163/125 | 94 | 28/26 | 4.89e-3/3.25e-5 | 859 |
| *COVID Serum* | 384 | 108/95 | 20 | 7/7 | 6.20e-1/4.52e-1 | 301 |

For each comparison, we report the number of differentially expressed (DE) metabolites, the number of enzyme-coding genes predicted by hypeR-GEM using both non-directional and directional mapping rules, and the number of DE genes/proteins from the paired omics layer. The non-directional and directional mapping strategies correspond to connecting metabolites through all reactions in which they participate or only through reactions in which they are produced, respectively (see “Materials and Methods”). We further evaluated the significance of the overlap between the predicted enzyme-coding genes and DE genes/proteins via Fisher test, with the background defined as the intersection of enzyme-coding genes represented in Human-GEM and the genes/proteins measured in the paired omics layer.

A dataset-level summary is visualized in Figure 3A, while Figure 3C summarizes the results by specimen category. Notably, 9 out of the 11 comparisons exhibited statistically significant overlap (p < 0.05) under at least one mapping rule, supporting the effectiveness of hypeR-GEM at the single-molecule level. Significant overlaps were observed across all specimen categories. Strong concordance was particularly evident in cell-line and tissue datasets – including EMT, serine starvation, kidney, and liver – where the sources of molecular signals are relatively homogeneous. Importantly, comparisons involving plasma-based and urine samples also demonstrated statistically significant overlap, which further validate the practical utility of hypeR-GEM.

**Figure 3.**
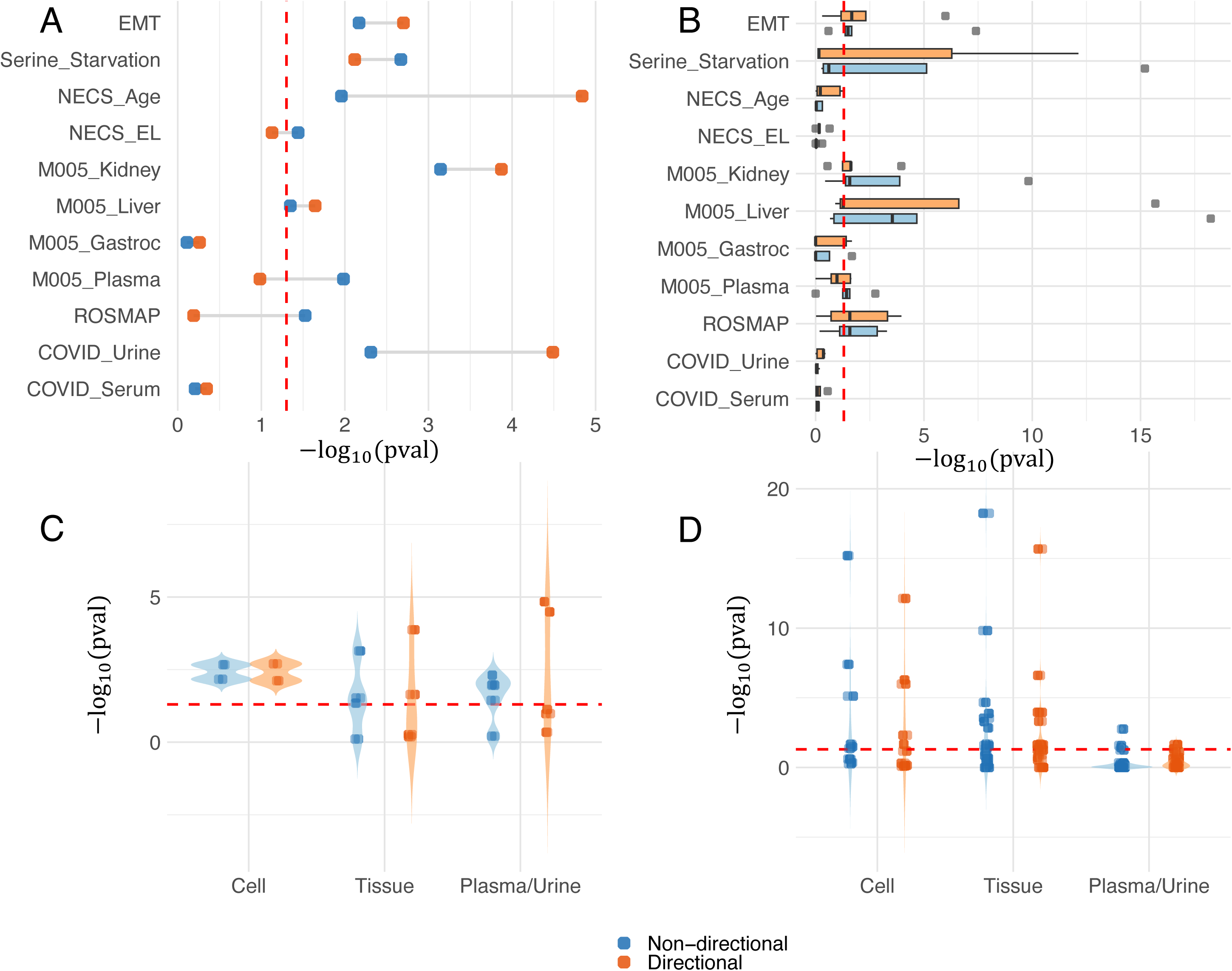
Single-molecule level and pathway level evaluation summary. The red dash line represents the α = 0.05 significance level. (A) Significance of the single-molecule level overlap between enzyme-coding genes predicted by hypeR-GEM and the corresponding proteomic or transcriptomic signatures for each dataset. (B) Significance of the pathway level overlap between enriched pathways identified by hypeR-GEM predicted enzyme-coding genes and those identified by annotation of the corresponding proteomic or transcriptomic signatures for each dataset. (C) Significance of the overlaps at the single-molecule level grouped by experimental model type. (D) Significance of the overlaps at the pathway level grouped by experimental model type.

#### Pathway level

Gene set ORA analysis was independently performed on 1) HypeR-GEM-mapped enzyme-coding genes, and 2) Corresponding proteomic or transcriptomic signatures. Pathway databases used included Hallmark^35^, Kyoto Encyclopedia of Genes and Genomes (KEGG)^36^, REACTOME^37^, Gene Ontology Biological Process (GOBP)^38^ and Gene Ontology Molecular Function (GOMF)^39^. We tested whether pathways enriched in signatures obtained from hypeR-GEM-mapped genes significantly overlapped with those obtained from the corresponding proteomic/transcriptomic data (see “Materials and Methods” for full details). Only comparisons where both omics layers yielded at least one enriched pathway were reported, see Table 2.

**Table 2.**
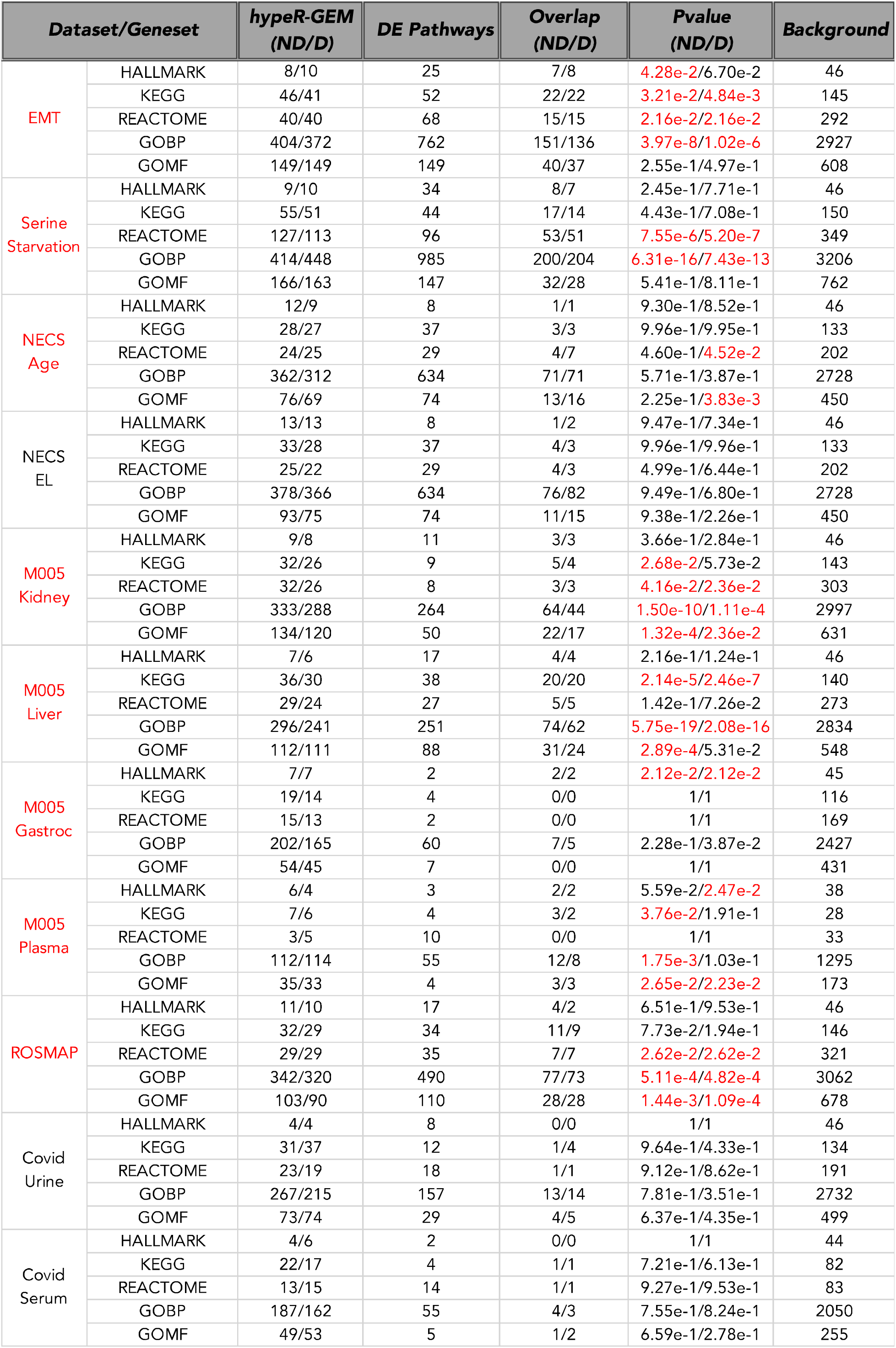
Evaluation of hypeR-GEM’s performance at the pathway level. *hypeR-GEM (ND/D)*: Number of enriched pathways obtained from hypeR-GEM mapped enzyme-coding genes based on non-directional (ND) and directional (D) mapping. *DE Pathways*: Number of enriched pathways obtained from transcriptomics/proteomics signatures. *Overlap (ND/D)*: The overlap between “hypeR-GEM” and “DE Pathways”. *Pvalue (ND/D)*: The p-value of the hypergeometric test. *Background*: The background parameter of the hypergeometric test, defined as the intersection of gene sets that are sufficiently represented in GEM and the corresponding proteomic or transcriptomic dataset.

For each comparison, we report the number of enriched pathways identified from hypeR-GEM predicted genes via both non-directional and directional mapping rules, as well as the number of enriched pathways derived from the DE genes/proteins in the paired omics layer. As in the single-molecule level evaluation, we applied a hypergeometric test to assess the statistical significance of the pathway-level overlap. The background for this test was defined as the set of pathways with sufficient representation in both Human-GEM and the corresponding transcriptomic or proteomic dataset.

A dataset-level summary is visualized in Figure 3B, while specimen-level results are summarized in Figure 3D. Notably, 7 out of the 9 comparisons that showed significant overlap at the single-molecule level also demonstrated statistically significant pathway-level overlap (p < 0.05) in at least one pathway database. Moreover, the M005 gastrocnemius muscle dataset – which did not show significant overlap at the single-molecule level – exhibited significant overlap in at least one pathway collection at the pathway level. In total, 8 out of 11 comparisons, spanning all specimen categories (cell, tissue, plasma, and urine), demonstrated statistically significant pathway-level concordance, further supporting the efficacy and generalizability of hypeR-GEM for functional interpretation of metabolomics data.

Global rank-based Kolmogorov-Smirnov tests were performed to assess whether the distributions of p-values were significantly skewed toward zero, with both the single-molecule and pathway level tests yielding highly significant results (Table 3).

**Table 3.** Global KS test. *Level*: Evaluation level. *Statistics*: The statistics of the KS test. *Pvalue*: The p-value of the KS test.

| **Level** | **Mapping** | **Statistics** | **Pvalue** |
| --- | --- | --- | --- |
| Single-molecule | Non-directional | 0.77 | 1.01E-07 |
|  | Directional | 0.62 | 5.90E-05 |
| Pathway | Non-directional | 0.33 | 8.42E-06 |
|  | Directional | 0.34 | 2.32E-06 |

## APPLICATION

We further demonstrate the practical utility of hypeR-GEM for multi-omics integration through a use-case using metabolomics data from the New England Centenarian Study (NECS). The dataset comprises 1,213 measured and 888 annotated metabolites. A signature of age-associated metabolites was identified by regressing each metabolite on age while controlling for sex, batch and years of education. This analysis identified 209 up-regulated and 84 down-regulated metabolites (adjusted p-value < 0.05). Among the up-regulated metabolites, 84 were represented in GEM and mapped to 307 and 250 enzyme-coding genes under the non-directional and directional mapping rules, respectively. Similarly, of the 84 down-regulated metabolites, 29 were represented in GEM and mapped to 274 and 190 enzyme-coding genes under the two mapping schemes. ORA-based enrichment analyses were performed both in metabolite and gene space, and the results are summarized in the three-layer Sankey diagram shown in Figures 4 (see “Materials and Methods” for full details), linking age-associated metabolites, enriched metabolite sets, and gene sets.

**Figure 4.**
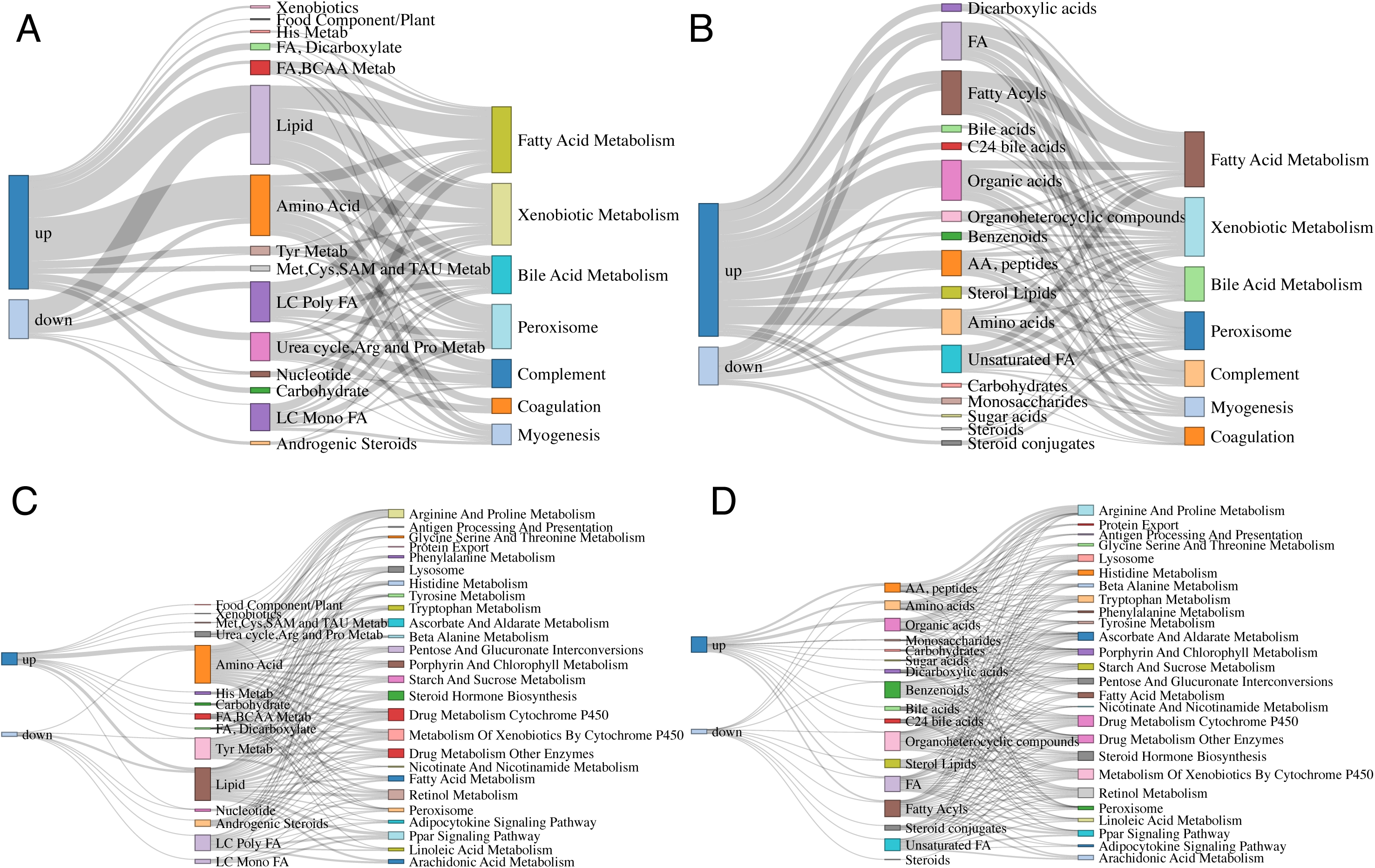
Metabolite- and gene-based annotation of age-associated signatures. Three-layer Sankey diagrams illustrating the connections between age-associated metabolites, enriched metabolite sets, and enriched gene sets (from left to right). Gene set ORA was performed using the HALLMARK and KEGG compendia. (A) Age-associated metabolites, Metabolon-annotated metabolite sets, and HALLMARK gene sets. (B) Age-associated metabolites, RefMet-annotated metabolite sets, and HALLMARK gene sets. (C) Age-associated metabolites, Metabolon-annotated metabolite sets, and KEGG gene sets. (D) Age-associated metabolites, RefMet-annotated metabolite sets, and KEGG gene sets.

Age-associated metabolites were significantly enriched in multiple lipid-related pathways across both annotation systems – namely, “Lipid” and “Long Chain Polyunsaturated Fatty Acids (LC Poly FA)”, “Long Chain Monounsaturated Fatty Acids (LC Mono FA)”, “Fatty Acids, Dicarboxylate (FA, Dicarboxylate)”, among others (Metabolon^40,41^; Figure 4A, 4C), as well as ““Fatty Acids (FA)”, “Unsaturated Fatty Acids (Unsaturated FA)”, “Bile acids”, “Fatty Acyls”, and “Sterol Lipid”, among others. (RefMet; Figure 4B, 4D). Notably, these metabolite-level lipid-based pathways are also linked to gene-level lipid-associated pathways enriched in hypeR-GEM predicted genes, including “Bile Acid Metabolism”, “Fatty Acid Metabolism”, “Retinol Metabolism”, “Steroid Hormones Biosynthesis”, “Tryptophan Metabolism”, and “Lysosome” pathways (from Hallmark and KEGG compendia). These cross-omics connections indicate coordinated lipid-associated enzymatic and cellular processes that span both the metabolite and gene layers^31,42–47^ (Table S12).

Such findings suggest that aging is accompanied by systemic reprogramming of lipid metabolism, and that perturbations in lipid pathways may propagate across omics layers. For instance, alternations in polyunsaturated fatty acids and bile acid metabolism may reflect altered membrane dynamics^48–50^, proteostatis^51,52^ and lipid signaling^52,53^, all of which are known to be associated with hallmarks of aging, including mitochondrial dysfunction, nutrient sensing, and chronic inflammation^54^.

Interestingly, the gene-level annotation highlighted, among others, enrichment of Retinol Metabolism in younger subjects (Table S12, KEGG). This was not explicitly captured at the metabolite level, and reflected decreased abundance with age of a varied list of metabolites not directly connected to Retinol metabolism, including Androsterone 3-glucuronide, Bilirubin, Thyroxine (T4), fatty acids that function as substrates for enzymatic oxidation pathways (arachidonic acid and linoleic acid)^55,56^, beside Retinol itself. The associated genes driving this enrichment, however, included an interconnected system of enzymes, ADH, CYP, UGT, and fatty acid/retinol metabolizing proteins (see Table S12), which enables the processing of hormones, vitamins, and lipid signaling molecules by a combination of oxidation, reduction, and conjugation^57–62^. The observed decrease in abundance of these metabolites and alteration in enzyme function with age likely contributes to impaired metabolic flexibility and increased oxidative stress, processes closely linked to aging and its associated functional decline.

Taken together, these findings align with the well-established role of lipid metabolism in aging and demonstrate the ability of hypeR-GEM to uncover biologically validated mechanisms, highlighting its effectiveness in real-world settings and underscoring its value for elucidating mechanistic insights into complex phenotypes such as aging.

## DISCUSSION

Enrichment analysis in metabolomics is often hindered by the limited availability of analytical tools, the absence of a comprehensive repository of molecular signatures based on prior knowledge, and the lack of well-defined associations between metabolites and genes. To address these challenges, we have developed hypeR-GEM, an R package that leverages gene–reaction–metabolite relationships from genome-scale metabolic models (GEMs) to enable gene set enrichment analysis on metabolomics data.

We evaluated hypeR-GEM across 11 phenotypes from 10 paired metabolomics-proteomics and metabolomics-transcriptomics datasets. At the single-molecule level, 9 out of 11 comparisons showed statistically significant overlap between hypeR-GEM–predicted enzyme-coding genes and differentially expressed genes/proteins from the paired omics layer. Furthermore, 7 out of these 9 also demonstrated significant pathway-level concordance (p < 0.05) in at least one gene set database (Hallmark, KEGG, REACTOME, GOBP, and GOMF).

We wish to emphasize that our evaluation relies on a stringent assumption – that genes/proteins associated with differentially expressed metabolites should also be differentially expressed – which is likely not a necessary condition. The concordance observed under this stringent assumption thus further underscores the utility and robustness of hypeR-GEM. Beyond assessing the statistical significance of the overlap, future evaluations could assess hypeR-GEM’s ability to recover literature-curated disease-associated signatures or pathways. Moreover, researchers can combine the mapped enzyme-coding genes with their domain-specific knowledge to formulate testable hypotheses.

To further demonstrate its practical utility, we applied hypeR-GEM to age-associated metabolic signatures from the NECS study. This analysis revealed consistent enrichment in lipid-related pathways, including Lipid Metabolism, Fatty Acid Metabolism, Retinol, Tryptophan, Lysosome, and Bile Acid Metabolism. These results are consistent with the well-established role of lipid metabolism in aging, which contributes to mitochondrial dysfunction, dysregulated nutrient sensing, and chronic inflammation^42,43,46,47,52^. These findings validate hypeR-GEM’s utility in analyzing complex biological mechanisms.

HypeR-GEM is an open-source R package released under the GNU General Public License, and it is available for Linux, macOS, and Windows on GitHub. Comprehensive documentation, tutorial vignettes, and workflow examples are hosted at https://github.com/montilab/hypeR-GEM. In summary, hypeR-GEM offers a user-friendly tool designed to leverage the advantages of gene set enrichment analysis, thereby facilitating the interpretation of complex results from metabolomics studies.

## MATERIALS AND METHODS

### Algorithm Framework

The core of hypeR-GEM is a reaction-centric mapping algorithm that systematically links metabolites to enzyme-coding genes based on the biochemical relationships encoded within genome-scale metabolic models (GEMs) (Figure 1). Given an input metabolic signature, a set of metabolites annotated using the RefMet nomenclature, hypeR-GEM identifies all metabolic reactions in which each metabolite participates as either reactant or product. For reversible reactions, metabolites are treated as both reactants and products. For each reaction (excluding exchange reactions and a subset of transport reactions), the associated enzyme-coding genes are then mapped to the metabolites according to two configurable rules: a *non-directional* mapping that considers all reactions involving the metabolite, and a *directional* mapping that restricts associations to reactions producing the metabolite based on the stoichiometry encoded in the model.

To quantify the specificity of each mapped gene’s association to the input signature, hypeR-GEM computes a *gene-specificity score* based on the hypergeometric distribution. This test statistically assesses the overlap between signature metabolites and those linked to each gene via GEM reactions. The resulting gene-specificity scores are used as weights in a hypergeometric test for gene set overrepresentation analysis, thereby prioritizing genes with highly specific connections to the input metabolite signature.

### Human-GEM

In this study, we utilized the open-source human genome-scale metabolic model (Human-GEM V1.17.0) for analysis and validation. Human-GEM encompasses 13,026 metabolic reactions and 8,365 metabolites across various compartments (e.g. cytosol, extracellular, etc.) and includes 3,068 enzyme-coding genes. The model is accessible at https://github.com/SysBioChalmers/Human-GEM. Notably, we used the model solely as a curated knowledge base to obtain gene–reaction–metabolite associations and to map metabolites to enzyme-coding genes.

In addition to the human model, the information of genome-scale metabolic models (GEMs) for five major model organisms – Mouse, Rat, Zebrafish, Fruitfly, and Worm – was also extracted and integrated into hypeR-GEM. Further details regarding these models can be found in Wang et al. (2021)^63^.

### Reaction-based metabolite-to-gene mapping

A prototypical reaction in GEMs comprises three fundamental components: reactants, representing a set of metabolites consumed by the reaction; products, comprising a set of metabolites produced by the reaction; and enzymes, which are responsible for catalyzing the reaction^64,65^. This gene–reaction–metabolite relationship provides a natural connection between metabolites and enzyme-coding genes. Specifically, for each metabolite of interest, we identified the reactions in which the metabolite participates, determined its roles (as reactant or product), and identified the genes encoding the corresponding enzymes catalyzing these reactions. Because hypeR-GEM uses GEMs solely as a curated repository of gene–reaction–metabolite relationships rather than for predicting reaction activity, we do not evaluate the Boolean gene–protein–reaction (GPR) rules that specify isoenzymes or multi-subunit enzyme complexes. Instead, all genes listed in the GPR annotation for a reaction are included as associated enzyme-coding genes. We then established two mapping rules: a non-directional rule, mapping the metabolite to the enzyme-coding genes in a reaction if the metabolite serves as either a reactant or a product of that reaction; and a directional rule, mapping the metabolite to the genes only if the metabolite acts as a product of the reaction. By iteratively applying these mapping rules to each metabolite within the signatures, the signatures can be transformed from metabolite-space to gene-space.

### Gene Specificity

Enzyme-coding genes are present in all living cells, where they catalyze and regulate reactions essential to the existence of the living system^66,67^. One important feature of enzymes is called reaction specificity, which depends on the shape of their active site^68–70^. Non-specific enzymes can catalyze a wide range of reactions, resulting in promiscuous gene–reaction–metabolite connections, and consequently, less specific associations with a given metabolic signature. To address this, we compute a gene-specificity score based on the hypergeometric distribution to evaluate the association specificity of each enzyme-coding gene (see Figure 5). The computed score measures the statistical significance of the overlap between the metabolites associated with the gene across the entire GEM and those within the metabolite signature (Figure 5).

**Figure 5.**
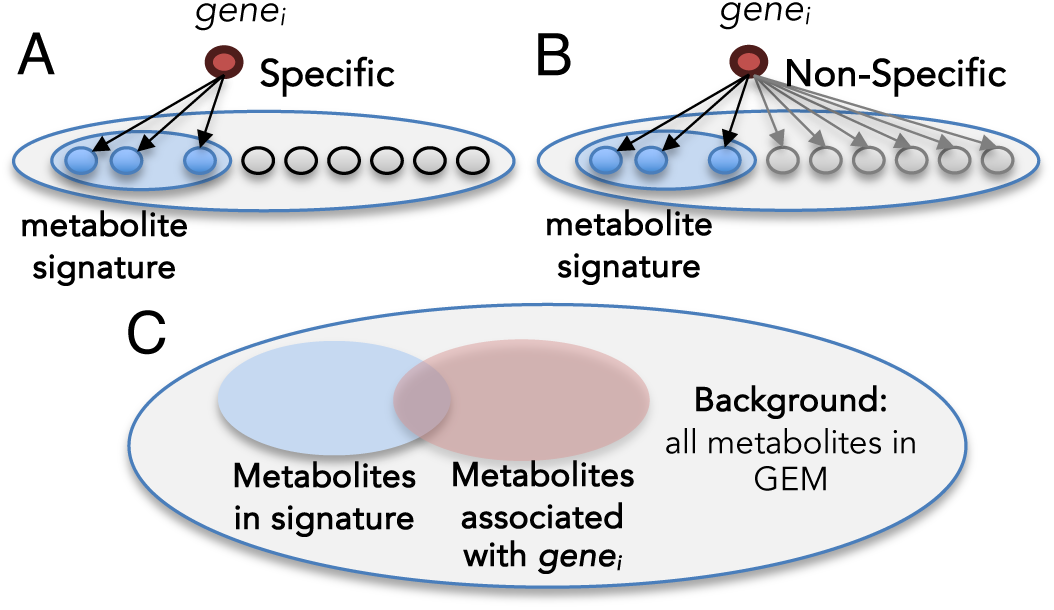
Illustration of gene-specificity. (A) Enzyme-coding genes that catalyze a limited number of reactions exhibit higher specificity in their association with metabolic signatures. (B) In contrast, non-specific enzymes, which catalyze more reactions, lead to more promiscuous gene–reaction–metabolite connections. (C) gene-specificity hypergeometric test quantifies the specificity of a given gene by assessing the statistical significance of the overlap between metabolites associated with the gene and those within a specific metabolic signature. The background for this test is defined as the total number of metabolites in the Human-GEM.

For each gene *g_i_*, let *M* denote he total number of metabolites represented in the GEM, *S* the number of metabolites in the metabolic signature, *A_i_* the number of metabolites associated with *g_i_* and *a_i_* the number of those metabolites that overlap with the signature. The significance score *s_i_* is defined as the upper-tail hypergeometric probability of observing at least *a_i_* overlapping metabolites by chance:

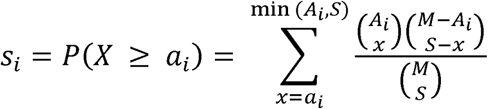

Smaller values of *s_i_* indicate more specific association. To incorporate this score into downstream enrichment analysis, each gene is assigned a weight *w_i_* ∈ [0,1] derived from a sigmoid transformation of its significance score:

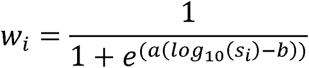

The sigmoid function is monotonic and bounded within [0, 1], assigning larger weights to genes with higher specificity. The parameter *b* determines the inflection point of the curve (the *half-point*, where *w_i_* = 0.5), allowing users to control the weight threshold, while *a* > 0 regulates the steepness of the curve.

### Weighted Hypergeometric test

The standard hypergeometric test employed in ORA assesses whether a set of genes exhibits an over-representation of genes belonging to a specific category (e.g., pathway or ontology term), assuming that each gene contributes equally to the enrichment^71,72^. However, as shown in the “Gene Specificity” section, GEM-mapped enzyme-coding genes vary in their association specificity with the input metabolic signature. To incorporate this information into enrichment analysis, we extend the standard test to a weighted hypergeometric test that uses the gene-specificity weights *w_i_*.

Formally, the probability of finding *X* ≥ *k* genes in a particular gene set or pathway is given by:

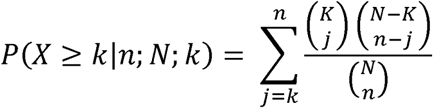

In the conventional formulation, N represents the size of background, n is the number of GEM-mapped enzyme-coding genes, K is the number of genes in each gene set or pathway, and k is the size of overlap.

In the weighted formulation, we first compute the continuous, weight-adjusted counts:

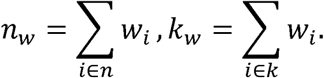

Because the hypergeometric test requires integer inputs, these continuous counts are rounded to the nearest integer using the standard notation

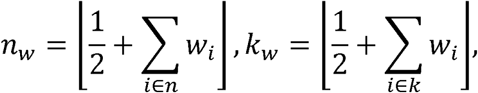

where ⌊·⌋ denotes the floor operator.

By substituting *n_w_* and *k_w_* into the hypergeometric test formula, the weighted version effectively down-weights highly promiscuous genes that exhibit low specificity, thereby mitigating noise, reducing spurious enrichments, and improving the biological relevance of the enrichment results.

### Implementation

HypeR-GEM is implemented entirely in R^73^. The core function, *signature2gene()*, accepts one or more input metabolic signatures and maps them to enzyme-coding genes using reaction-level information from genome-scale metabolic models (GEMs). Input metabolites must be annotated using the RefMet nomenclature ^74^. The output is a structured list object containing metadata required for downstream gene set ORA analysis, including Ensembl IDs ^75^, associated metabolites, the number of catalyzed reactions, and hypergeometric test statistics quantifying each the signature-specificity of each mapped gene.

HypeR-GEM provides a modular workflow that facilitates not only the mapping of metabolites to genes, but also downstream gene set ORA (function *enrichment()*), visualization (functions *enrichment_plot(), rctbles(), sankey_plot()*), and generation of interactive reports (function *hypGEM_to_excel()*). Users may apply their preferred statistical analysis pipelines to metabolomics data to derive statistically significant metabolite signatures, which can then be analyzed and interpreted in the gene space using hypeR-GEM. Output can be easily exported to Excel-compatible format, enabling reproducible and shareable reports.

### Dataset overview

To evaluate the performance of hypeR-GEM, we utilized paired metabolomics–proteomics and metabolomics–transcriptomics datasets from ten independent studies, covering four specimen categories and conducting differential analyses across eleven phenotypes (see Table 4). Results from the differential analyses and GEM-mapping are provided in Supplementary Data S1 to S11, with only molecules meeting the significance threshold (adjusted *p*-value < 0.05) were reported.

**Table 4.** Overview of paired datasets used for method evaluation. *Pair:* The paired omics layers. *Metabolites*: The number of metabolites. *Genes/Proteins*: The number of proteins. *Phenotype*: The phenotype used for differential analysis. *Specimen:* The biological specimen from which the samples were derived. *EL*: Extreme longevity. *SnellWT*: Snell wild type mice. *SnellDW*: Snell dwarf mice. *NCI*: No cognitive impairment. AD: Alzheimer dementia.

| **Dataset** | **Pair** | **Metabolites** | **Genes/Proteins** | **Phenotype** | **Specimen** |
| --- | --- | --- | --- | --- | --- |
| *EMT* | Metabolomics+<br>Proteomics | 8325 | 6426 | Epithelial vs.<br>Mesenchymal | Cell |
| *Serine Starvation* | Metabolomics+<br>Transcriptomics | 318 | 15393 | Serine-restriction vs.<br>Control | Cell |
| *NECS Age* | Metabolomics+<br>Proteomics | 1052 | 4783 | Age | Plasma |
| *NECS EL* | Metabolomics+<br>Proteomics | 1052 | 4783 | EL vs. Not | Plasma |
| *M005 Kidney* | Metabolomics+<br>Proteomics | 759 | 5358 | SnellDW vs.<br>SnellWT | Tissue |
| *M005 Liver* | Metabolomics+<br>Proteomics | 509 | 4019 | SnellDW vs.<br>SnellWT | Tissue |
| *M005 Gatroc* | Metabolomics+<br>Proteomics | 947 | 903 | SnellDW vs.<br>SnellWT | Tissue |
| *M005 Plasma* | Metabolomics+<br>Proteomics | 971 | 2384 | SnellDW vs.<br>SnellWT | Plasma |
| *ROSMAP* | Metabolomics+<br>Proteomics | 1055 | 8817 | NCI vs. AD | Tissue |
| *COVID Urine* | Metabolomics+<br>Proteomics | 1033 | 4003 | COVID-19 vs.<br>Control | Urine |
| *COVID Serum* | Metabolomics+<br>Proteomics | 903 | 1494 | COVID-19 vs.<br>Control | Plasma |

### TGFβ-induced epithelial-to-mesenchymal transition in MCF10A cells

Epithelial-to-mesenchymal transition (EMT) is a biological process in which polarized epithelial (E) cells lose their specialized features, pass through intermediate hybrid states (E/M), and acquire mesenchymal (M) characteristics. This transition regulates cell plasticity in contexts such as embryonic development, wound healing, fibrosis, and cancer^28,76^. For this study, we analyzed paired metabolomics and proteomics datasets obtained from MCF10A cells exposed to TGFβ across ten time points, each with three biological replicates. Following Paul *et al.* (2023)^23^, cells from the first three time points were classified as epithelial (E), those from the last four time points as mesenchymal (M), and the remaining time points as intermediate (E/M). The metabolomics dataset comprised 8,325 metabolites, while the proteomics dataset included 6,426 proteins. Differential analyses were performed by comparing cells in the epithelial (E) state with those in the mesenchymal (M) state, using a mixed-effects model that considered biological replicates as a random effect (see Table 4). The raw data are publicly available through ProteomeXchange (Proteomics: accession PXD031071) and the National Metabolomics Data Repository (Metabolomics: PR001174), as described in Paul *et al.* (2023)^28^.

### Serine starvation in HSC3 cells

Oral squamous cell carcinoma is an aggressive malignancy with limited therapeutic options. Emerging evidence highlights the critical role of serine, a non-essential amino acid, in regulating cell fate decisions, and demonstrates that serine restriction can suppress tumor progression across several cancer models. In this study, we analyzed paired metabolomics and transcriptomics datasets comprised 6 samples obtained from HSC3 cells cultured under serine-restricted versus control conditions (3 samples per condition). The metabolomics dataset comprised 318 metabolites, while the transcriptomics dataset comprised 15,393 transcripts (see Table 4). The RNA-seq data are available in the Gene Expression Omnibus (GEO; accession GSE286464), as reported by Jankowski *et al.* (2025). (manuscript in preparation).

### The New England Centenarian Study (NECS)

The New England Centenarian Study (NECS) provides a unique resource for the investigation of the biological factors contributing to extreme human longevity (EL)^30^. We leveraged previously generated datasets obtained from serum samples subjected to proteomics^30^ and untargeted metabolomics (manuscript in preparation) profiling. The metabolomics dataset included 1,052 metabolites that passed quality control across 213 subjects, comprising 70 centenarians, 80 offspring and 63 control. The proteomics dataset^30^ included 4,783 proteins across 224 subjects comprising 77 centenarians, 82 offspring, and 65 control, with 205 subjects in common between the two datasets. These were subsequently analyzed to identify age- and EL-associated molecular signatures (see Table 4).

### M005

The M005 dataset originates from the NIA Longevity Consortium’s lifespan-extension studies in genetically heterogeneous UM-HET3 mice^31^. It comprises paired metabolomic and proteomic profiles from kidney, liver, gastrocnemius muscle (Gastroc), and plasma specimens, providing a systems-level resource for investigating molecular mechanisms of aging and longevity. Differential analyses were performed across all four specimen types by comparing 17 Snell dwarf (SnellDW) mice, which carry a loss-of-function mutation in the *Pit1* gene leading to reduced body size, delayed sexual maturation, and extended lifespan^31^, with 18 wild-type (SnellWT) controls. The numbers of metabolites and proteins in the metabolomics and proteomics datasets are summarized in Table 4.

### ROSMAP

The Religious Orders Study (ROS), initiated in 1994, and the Rush Memory and Aging Project (MAP), initiated in 1997, are longitudinal cohort studies collectively referred to as ROSMAP^32,33^. These cohorts investigate risk factors for cognitive decline, Alzheimer’s disease, and related health outcomes. The metabolomics dataset^32^, derived from brain tissue, included 1055 metabolites that passed the quality control across 521 subjects, comprising 155 individuals with no cognitive impairment (NCI), 197 individuals with Alzheimer’s dementia (AD), and the remaining subjects have varying degrees or alternative causes of cognitive impairment. The proteomics dataset^77–80^, also generated from brain tissue, included 8,817 proteins across 400 subjects, comprising 168 individuals with no cognitive impairment (NCI), 109 individuals with Alzheimer’s dementia (AD), and the remaining subjects have varying degrees or alternative causes of the cognitive impairment. We performed differential analyses comparing individuals with no cognitive impairment (NCI) to those with Alzheimer’s dementia (AD) (see Table 4).

### COVID-19

This dataset derives from a clinical study of hospitalized COVID-19 patients in Taizhou, China, which systematically profiled the urinary and serum proteomes and metabolomes of 71 patients with COVID-19 together with 27 healthy controls^34^. Differentially expressed metabolites and proteins were identified by comparing COVID-19 disease with healthy individuals. We utilized the publicly available COVID-19–associated metabolite and protein signatures reported in this study^34^ (see Table 4).

### Differential analysis and single molecule-level evaluation

Differential analysis for each dataset (excluding COVID-19 datasets) was conducted using linear regression in R. For each paired dataset, differential analysis was performed independently on each omics layer to identify differentially expressed molecules, or signatures. For the COVID-19 datasets, we utilized publicly available signatures that had been previously identified^25^. The p-values of each molecule were adjusted using the Benjamini-Hochberg method, the statically significance was assessed using an adjusted *p*-value threshold of < 0.05. The resulting signatures from each omics were further classified as up- or down-regulated based on the sign of the estimated coefficient. Up- and down-regulated metabolites were then used separately as inputs for hypeR-GEM to obtain mapped enzyme-coding genes. The evaluation was carried out by performing a hypergeometric test to assess the overlap between the union of hypeR-GEM-mapped enzyme-coding genes and the union of up- and down-regulated proteins/genes obtained from the corresponding proteomics/transcriptomics data.

### Gene set enrichment analysis and pathway-level evaluation

Gene set enrichment analysis was conducted using the *hypeR*^81^ package in R on pathways from the Hallmark^35^, Kyoto Encyclopedia of Genes and Genomes (KEGG)^36^, REACTOME^37^, Gene Ontology Biological Process (GOBP)^38^ and Gene Ontology Molecular Function (GOMF)^39^. This analysis was conducted independently on the on the enzyme-coding genes predicted by hypeR-GEM and signatures derived from the corresponding proteomic/transcriptomic signatures, with a significant threshold of adjusted p-value < 0.05. The evaluation employs hypergeometric test to assess the overlap between the enriched pathways identified from hypeR-GEM-mapped genes and those obtained from proteomics/transcriptomics signatures.

### Application to Age-Associated Metabolic Signatures in NECS

We analyzed age-associated metabolic signatures derived from the NECS metabolomics dataset and performed Metabolite Set Enrichment Analysis using pathway annotations from Metabolon, Inc.^40,41^ and Refmet classification system^74^, identifying metabolite-level enriched pathways based on an adjusted p-value threshold of ≤ 0.05. Additionally, we applied hypeR-GEM to map these age-associated metabolites to enzyme-coding genes, followed by Gene Set Enrichment Analysis using curated gene sets from Hallmark^35^ and Kyoto Encyclopedia of Genes and Genomes (KEGG)^36^, identifying enriched gene-level pathways (adjusted p-value < 0.05).

To integrate these complementary layers into a unified biological context, we constructed three-layer Sankey diagrams connecting differentially expressed metabolites, the enriched metabolite sets (MSets), and the enriched gene sets (GSets) derived from hypeR-GEM. Edge weights between the metabolite and Msets layers reflect the number of shared metabolites, while edges between the MSets and GSets layers reflect the overlap between hypeR-GEM–mapped genes and genes in each GSets. This framework enables the joint visualization of age-associated mechanisms across both metabolite and gene/protein spaces.

## Supporting information

Main Text Tables

## SUPPLEMENTARY DATA

Supplementary Data S1 to S11 contain the analysis results for the eleven phenotypes analyzed across ten datasets, covering four specimen categories. Only molecules that satisfied the significance threshold (adjusted p-value < 0.05) are reported. Each Excel file corresponds to a single phenotype and is organized into four sheets: (1) *Diffanal Proteomics* (differential analysis of proteomics), (2) *Diffanal Metabolomics* (differential analysis of metabolomics), (3) *GEM-mapped Non-directional* (enzyme-coding genes mapped via the non-directional rule), and (4) *GEM-mapped Directional* (enzyme-coding genes mapped via the directional rule).

Supplementary Data S12 contains the enrichment analysis of age-associated metabolite signatures from the New England Centenarian Study (NECS). The Excel file includes: (1) over-representation analyses of Metabolon-annotated and RefMet-annotated metabolite sets using age-associated metabolite signatures, and (2) over-representation analyses of the HALLMARK, KEGG, and REACTOME gene-set compendia using hypeR-GEM–mapped enzyme-coding genes derived from these metabolite signatures.

## DATA AND CODE AVAILABILITY

### EMT data

The unprocessed data are publicly available through ProteomeXchange (Proteomics: accession PXD031071) and the National Metabolomics Data Repository (Metabolomics: PR001174), as described in Paul *et al.* (2023)^28^

### Serine Starvation data

The RNA-seq data are available in the Gene Expression Omnibus (GEO; accession GSE286464), as reported by Jankowski *et al.* (2025)^29^. (waiting for metabolomics GSE #)

### ROSMAP data

Study data were provided by the Rush Alzheimer’s Disease Center, Rush University Medical Center, Chicago. Data collection was supported through funding by NIA grants U01AG46161(TMT proteomics), the Illinois Department of Public Health (ROSMAP). Additional phenotypic data can be requested at www.radc.rush.edu. Metabolomics data is provided by the Alzheimer’s Disease Metabolomics Consortium (ADMC) and funded wholly or in part by the following grants and supplements thereto: NIA R01AG046171, RF1AG051550, RF1AG057452, R01AG059093, RF1AG058942, U01AG061359, U19AG063744 and FNIH: #DAOU16AMPA awarded to Dr. Kaddurah-Daouk at Duke University in partnership with a large number of academic institutions. As such, the investigators within the ADMC, not listed specifically in this publication’s author’s list, provided data along with its pre-processing and prepared it for analysis, but did not participate in analysis or writing of this manuscript. A complete listing of ADMC investigators can be found at: https://sites.duke.edu/adnimetab/team/. The Metabolon datasets were generated at Metabolon and pre-processed by the ADMC.

### M005 data

The M005 proteomics and metabolomics data is available from the ELITE portal (https://doi.org/10.7303/syn69931740).

### COVID data

The unprocessed proteomics data are publicly available through ProteomeXchange (Proteomics: accession PXD030662). The COVID-19–associated metabolite and protein signatures reported by Bi *et al.* (2022)^34^ are available in the supplementary materials of that study.

### NECS data

NECS data will be available from the ELITE portal.

All analyses in this study were conducted using R, and the scripts are available from the GitHub repository: montilab/huang_hypeR.GEM_analysis:v1.0.0 (https://doi.org/10.5281/zenodo.17108624).

Any additional information required to reanalyze the data reported in this work is available from the lead contact upon request.

## ACKNOWLEDGMENTS

We wish to thank Noa Rappaport, Greg Tombline, Rich Miller, and Oliver Fiehn, for useful discussions on the design of the method and its evaluation. The results published here are in part based on data obtained from the AD Knowledge Portal (https://adknowledgeportal.org).

## FUNDING

This work was supported by the National Institutes of Health, NIA cooperative agreements UH3AG064704, U19AG023122-16, and Find the Cause Breast Cancer Foundation (findthecausebcf.org). The findings and conclusions presented in this paper are those of the author(s) and do not necessarily reflect the views of the NIH or the U.S. Department of Health and Human Services.

## AUTHOR CONTRIBUTIONS

Conceptualization, S.M. and Z.H.; data curation, Z.H. and S.M.; data analysis, Z.H. and S.M.; results interpretation, Z.H., S.M, P.S., D.S.; manuscript writing, original draft, Z.H. and S.M.; writing – review & editing, all authors.

## Notes

https://github.com/SysBioChalmers/Human-GEM

https://www.metabolomicsworkbench.org/data/index.php

https://www.ncbi.nlm.nih.gov/geo/

https://sites.duke.edu/adnimetab/team/

https://www.synapse.org/Synapse:syn69931740.draft/datasets/

### Competing Interest Statement

The authors have declared no competing interest.

